# Relation of seasonal birth pulses and maternal immunity with viral invasion and persistence: A case study of Hendra virus infection in a population of black flying foxes (*Pteropus alecto*)

**DOI:** 10.1101/582726

**Authors:** Jaewoon Jeong, Alison J. Peel, Raina K. Plowright, Olivier Restif, Hamish Mccallum

## Abstract

Increasing outbreaks of emerging infectious diseases, originating from wildlife, has intensified interest in understanding the dynamics of these diseases in their wildlife reservoir hosts. Until recently, the effect of seasonal birth pulses and subsequent waning of maternally derived antibodies on epidemics in a wild mammal population has received little attention and has remained obscure. In this study, we explore how population structure, influenced by seasonal breeding and maternally derived immunity, affects viral invasion and persistence, using a hypothetical system loosely based on Hendra virus infection in black flying foxes (*Pteropus alecto*). We used deterministic epidemic models to simulate transient epidemics, following viral introduction into an infection-free population, with a variety of timings within a year and different levels of pre-existing herd immunity. Moreover, we applied different levels of birth synchrony and different modelling methods of waning maternal immunity to examine the effect of birth pulses and maternally derived immunity, both individually and in combination. The presence of waning maternal immunity dispersed the supply time of susceptible individuals in seasonally breeding populations, hence diminishing the effect of birth pulse. Dampened epidemics, caused by waning maternal immunity, made viral invasion and persistence easier. This study enhanced our understanding of viral invasion, persistence, and timing of epidemics in wildlife populations.

## Introduction

Seasonal behaviour of wildlife hosts—including aggregation, reproduction, migration, and associated physiological variations—influences the transmission of infection in their populations [1]. Among the seasonal factors, breeding has been implicated as playing a role in determining pathogen persistence and epidemic patterns. Seasonal births result in seasonal variation in population size and proportion of susceptible hosts, affecting infection of the population [2]. Seasonal breeding could result in a population requiring a much larger size to maintain a pathogen [3]. The frequency of birth pulses within a year has also been suggested as a major factor determining viral persistence [4]. Moreover, seasonal breeding can drive the timing of recurrent epidemics in a wildlife population [5].

Births from susceptible mothers and immigration of susceptible individuals are not the only sources of virus-susceptible individuals. While newborns of immune mothers can obtain protection against infection via maternally-derived antibodies (MatAb), this protection wanes over a certain period [6], causing these individuals to enter the susceptible pool as juveniles. The proportion of immune mothers that confer waning passive immunity to newborns could, therefore, significantly modify the supply of susceptible hosts. The supply of susceptible hosts from these two sources (from births to susceptible mothers and via loss of MatAb [7]) interact with each other to determine whether viral introduction into a population would result in viral invasion or persistence. The presence of MatAb in seasonally breeding wildlife is known to delay recurrent epidemics over a multi-year timescale [8]. However, the mechanism underlying the effect of this delay on the timing of epidemics within a year requires further investigation.

A modelling approach was established to describe the infectious period by an exponential distribution, which is a classical method to transfer individuals across infection stages [9] and is implicit in a simple differential equation SIR (susceptible-infectious-immune) model with constant rate parameters [10]. Although the exponential distribution has been used widely for computational ease, efforts towards finding more realistic methods to model the infectious period resulted in the development of gamma-distributed infectious periods [11]. Wearing, Rohani (12) addressed the significance of modelling the infectious and latency periods in predicting the impact of infectious diseases. Although maternal immune period has been modelled using a conventional exponential distribution, for both infectious and latency periods, the effect of this method on the model outcomes has been poorly studied. Therefore, it is worthwhile to explore the mechanism of generation of different epidemic modelling outcomes by an exponential and a gamma distribution. This would, in turn, help to determine whether the method of modelling waning maternal immunity is an important influence on model behaviour.

Bats (Order: *Chiroptera*) have been identified as natural reservoirs of many emerging infectious diseases of public health concern [13, 14]. They do not appear to suffer from many of these infections, especially those caused by viruses, despite the obvious exceptions of rabies and Tacaribe virus [15, 16]. However, the fatality rates attributable to bat viral diseases are often quite high in other mammalian species [13]. For example, Hendra virus has been detected in four species of flying foxes (*Pteropus alecto, P. conspicillatus, P. poliocephalus*, and *P. scapulatus*) in Australia [17].. Spillover from *P. alecto* and *P. conspicillatus* [18-20] to horses, and thereafter to humans, causes serious clinical symptoms and even death in both horses and humans [21]. Excretion of Hendra virus by flying foxes rarely results in spillover events; excreted virus has to overcome several barriers such as pathogen survival and spread in the environment, and domestic animal or human exposure [22, 23]. Surveillance of Hendra virus spillover events has revealed temporal patterns in the southern subtropics of eastern Australia between June and September [24]. Seasonal behaviour of reservoir hosts, such as seasonal breeding, needs to be investigated to examine how it can influence the seasonal characteristics of viral prevalence [3, 25]. Although seasonal factors such as birth pulses are not the only drivers of Hendra virus dynamics, and other factors (*Eucalyptus* phenology, bat movement, temperature, and nutrition) can also significantly influence Hendra virus spillover [22, 26], it would be valuable to understand the underlying mechanism of this influence, since Hendra virus spillover is known to result from a variety of processes with many interacting factors involved [23].

Using a system loosely based on Hendra virus infection in a population of *P. alecto*, we investigate the effect of seasonal birth pulses and waning MatAb on viral invasion and persistence in a certain population. Moreover, we explore the hypothesis that seasonal birth pulses and waning MatAb can facilitate the seasonal clustering of Hendra virus epidemics in *P. alecto*. We deterministically model a hypothetical population of *P. alecto* with two varying conditions of herd immunity (proportion of recovered individuals in the population at the time of viral introduction) and days from birth pulse to viral introduction (days from the last birth pulse before viral introduction until viral introduction) in every model.

## Methods

### Model structure

We simulated the Hendra virus dynamics in *P. alecto* using compartmental deterministic models, which were framed using ordinary differential equations (ODEs) and numerically integrated using the *deSolve* package [27] in R [28]. Profiles of Hendra virus prevalence and seroprevalence in flying foxes may be generated by at least three different underlying mechanisms: susceptible-infectious-recovered (SIR), susceptible-infectious-recovered-susceptible (SIRS), and susceptible-infectious-latent-infectious (SILI) [29]. Among the three scenarios, we use assume the SIR here, but recognize that other disease processes, such as SILI, are highly likely. The purpose of this study was to explore the effect of maternally derived immunity on viral prevalence; therefore, factors such as loss of acquired immunity and latent infection were excluded in our models to increase modelling transparency and to ease the interpretation of modelling results. In addition to the SIR model, an MSIR model was built by adding a maternally immune (M) compartment to the existing SIR model. Maternally immune (M) newborns become susceptible at the rate δ (Table 1). Susceptible (S) bats become infected at the rate βSI (assuming density-dependent transmission). Bats are infectious (I) before recovery at the rate γ. Based on the SIR model, we assumed that once recovered (R), bats remain immune for the rest of their life (modelled in [30] along with the scenario of latency and reactivation). Hendra virus infection is not known to not cause clinical disease in its reservoir hosts (flying foxes) [31]; therefore, infected bats are assumed to have the same mortality rate as susceptible and recovered bats. The models were simulated with annual birth pulses for eight years and run with a daily time step.

**Table 1.** Model parameters.

| Parameter | Symbol | Value | Unit | Reference |
| --- | --- | --- | --- | --- |
| Transmission rate | $\beta$ | 0.0000476 | Per capita per day | [30] |
| Recovery rate | $\gamma$ | 1/7 | Per capita per day | [30] |
| Mortality rate | $\mu$ | 1/7 | Per capita per year | [32] |
| Maternally-derived immunity losing rate | $\delta$ | 1/255 | Per capita per day | [33] |
| Ageing rate | $\alpha$ | 1/2 | Per capita per year | [34] |
| Scalar to control birth rate | $\kappa$ | 0.00159 | Per adult per day | This study |
| Low, medium and high birth pulse synchronies | $s$ | 0.013, 1.3, and 130 | - | [3] |
| Timing of birth pulse | $\phi$ | 1/2 | - | [32] |
| Initial herd immunity |  | 0.55~0.85 | - |  |
| Days from the last birth pulse to viral introduction |  | 0~365 | day |  |
| Colony size | $N$ | 10000 | capita | [30] |
| Average proportion of juvenile bats in total bats | $\eta$ | 0.23 | - | [32] |
| Gamma-distribution parameter | $g$ | 8 | | [33] |

Simulation of seasonal birth pulses and maternally-derived immunity required an age-structured model of bat population. The age-structure consisted of sexually immature juvenile bats (denoted by the subscript *i*) and sexually mature adult bats (subscript *m*). In the MSIR model, juveniles became adults, which could breed two years after their birth (at the rate ε) [34]. The proportion of juveniles in the total population (η) was, on average, 0.24 [32]. Four epidemic compartments for the two age groups led to a total of eight stages (see Supplementary Equations S1 and S2). Although maternally immune adults are not expected to exist in nature, we included this stage for modelling consistency. The exponential distribution of periods of maternally-derived immunity means that some individuals stay in the maternally immune compartment longer than in any period between birth and adulthood. With the parameters used, only few maternally immune adults remained, and hence had an insignificant effect on the results. We assumed an age-independent annual mortality rate (μ) of 16 % [32]. Mortality rate (μ) and birth rate were independent of population density and were so chosen that the population size remained constant. Bats born to immune female adults (*R*_*m*_) and to maternally immune female adults (*M*_*m*_) were assumed to be maternally immune (*M*_*i*_) in the MSIR model, whereas all newborns were assumed to be susceptible (*S*_*i*_) in the SIR model. The initial numbers in each compartment are described in Supplementary Text S1 and Supplementary Table S1.

Seasonally pulsed births were modelled with a periodic Gaussian function (PGF) [3, 4]: 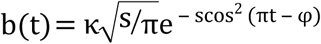, where κ controls the magnitude, s determines synchrony, and φ determines the timing of birth pulse. This function allows births to occur exclusively in a certain period within a year, with none outside this period [3, 4]. The scaling parameter (κ) was used so that the total population size would be stable inter-annually. *P. alecto* shows different seasonal birth patterns, depending on latitude [32]. In southeast Queensland and Northern NSW, where most Hendra virus spillover events have occurred till date, the offspring of *P. alecto* are known to be born in October and November, which is closely aligned to that of *P. poliocephalus* [32, 35]. With reference to the seasonal birth pulse of *P. poliocephalus*, we set s = 130, so that 95 % of annual births are concentrated within a one-month period [3] (see Supplementary Fig. S1 for annual fluctuation of the number of individuals). In addition to high levels of synchrony (s = 130), we also modelled low (s = 0.013) and medium (s = 1.3) levels of birth synchrony for comparison.

Flying fox colonies are patchily distributed, and individuals move among colonies. This movement may introduce Hendra virus into infection-free colonies, which may thereafter trigger transient epidemics (causing metapopulation dynamics of the virus within the host populations [30]). As the natural reservoir hosts of Hendra virus, flying fox colonies are highly likely to have been previously exposed to the virus [30], and so colonies are not necessarily immunologically naïve at the timing of viral introduction. Partial immunity of a population affects the duration and size of an epidemic caused by the viral introduction [36]. We therefore designed models to simulate Hendra virus dynamics in a colony of flying foxes within this broader context of a metapopulation in eastern Australia, assuming that a proportion of bats in a colony had previously been exposed to Hendra virus, and hence, was partially immune.

Although the transmission mode of Hendra virus among flying foxes is uncertain, a combination of frequency and density dependence has been hypothesised [24]. This study assumed density-dependent transmission, affected by population size, rather than frequency-dependent transmission, which remains unaffected by population size. Serological surveys of wild flying foxes showed an average proportion of susceptible individuals of 60 % [37, 38], and by with this finding, Plowright, Foley (30) estimated the mean values of a Hendra virus transmission rate (β) of 0.0000476 and recovery rate (γ) of 1/7 days^-1^, which we used in our models (Table 1). The loss rate of maternally derived immunity (δ) was assumed as 1/255 day^-1^, based on the data from an earlier experimental study on eight newborn *P. alecto* (born to seropositive bats) [33].

### Gamma-distributed periods of maternally-derived immunity

The MSIR model with exponentially distributed waning periods of MatAb (hereafter referred to as exponential MSIR model) assumed that the number of maternally immune hosts decreased exponentially since birth. In addition to the exponential MSIR model, an MSIR model with gamma-distributed waning periods of MatAb (hereafter referred to as gamma MSIR model) was simulated. This model used a gamma distribution in transferring maternally immune juveniles to susceptible juveniles. Gamma-distributed waning periods of MatAb were modelled by dividing the maternally immune stage into multiple (g) sub-stages, where gamma-distributed waning periods of MatAb had a mean of g and variance of 1/g [12] (see Supplementary Equation S3). As the gamma distribution parameter (g) increased, the rate of loss of maternally immune individuals shifted from an exponential decay to a fixed duration of maternal immunity. Epstein, Baker (33) determined the mean duration of maternal immunity (255 days) and half-lives for serum antibody against Hendra virus in twelve pups of *P. alecto*. Using the half-lives, we calculated the duration of maternal immunity in each pup so that the mean duration would be 255 days. We then modelled the gamma-distributed maternally immune durations with a variety of gamma distribution parameters and found that the mean duration was the most closely achieved with g = 8. Thus, we used the gamma distribution parameter (g = 8) for our models (Supplementary Fig. S2). This study included nine models: low, medium, and high levels of birth synchrony (s = 0.013, 1.3, and 130) in each of the three different modelling methods in terms of waning of MatAb: SIR model, exponential MSIR model, and gamma MSIR model.

### Pseudo-extinction

Viral persistence, which is an essentially stochastic process, should normally be analysed using stochastic models [39]. Nevertheless, deterministic models with a cut-off to determine viral persistence may provide valuable insights. We used *pseudo-extinction*, in which the number of infectious bats dropping below a cut-off was assumed to be a proxy for viral extinction in a population. If the sum of infectious juveniles (*I*_*i*_) and infectious adults (*I*_*m*_) dropped below one, in the population we assumed that the infection failed to persist, and hence, the simulations were stopped. We applied varying conditions of herd immunity and days from birth pulse to viral introduction, since herd immunity could be an important factor in determining the magnitude and duration of an epidemic [30] and days from birth pulse to viral introduction might affect viral persistence in seasonally breeding wildlife [3]. We used deterministic models over stochastic models, since the former were faster so that it was appropriate to use when we simulate with a wide ranges of herd immunity and timing of viral introduction. Since viral extinction is likely to occur in the first few post-epidemic troughs [40], the simulation period was limited to six years, which was considered to be long enough to include the first few epidemics and the following troughs.

## Results

Epidemics, caused by the introduction of an infectious bat into a bat population having a wide range of immunities, but no current infections, were simulated with the SIR model, exponential MSIR model, and gamma MSIR model. The predominant effect of herd immunity on epidemics, compared to the effect of birth synchrony and maternally-derived immunity, made the difference of epidemics insignificant in all three models at herd immunity < 0.55. However, differences occurred when herd immunity was similar to the herd immunity of an endemic equilibrium state (approximately between 0.55 and 0.7).

### Viral invasion

High birth synchrony (s=130) enhanced the importance of the effect of viral introduction time over low birth synchrony (s=1.3) (**Error! Reference source not found.**). With high birth synchrony (s=130), epidemics occurred at a broader range and higher levels of herd immunity. In particular, viral introduction shortly before a birth pulse (Fig. 1, DB >300), was found to result in the highest magnitude of epidemics. Compared to the SIR model, the addition of MatAb in the exponential MSIR model resulted in more uniform steady magnitude of epidemics across the timings of viral introduction at high birth synchrony (s=130), but higher maximum epidemic magnitudes at low and medium birth pulse synchrony (s=13). Compared to the exponential MSIR model, the gamma MSIR model showed even more uniform epidemic magnitudes across the timings of viral introduction (at high birth synchrony), with little difference observed at low and medium synchrony. Waning MatAb therefore appeared to reduce the effect of a highly seasonal birth pulse by dispersing the timing of supply of susceptible individuals, and this effect was most pronounced in the gamma MSIR model compared to the exponential MSIR.

**Fig. 1.**
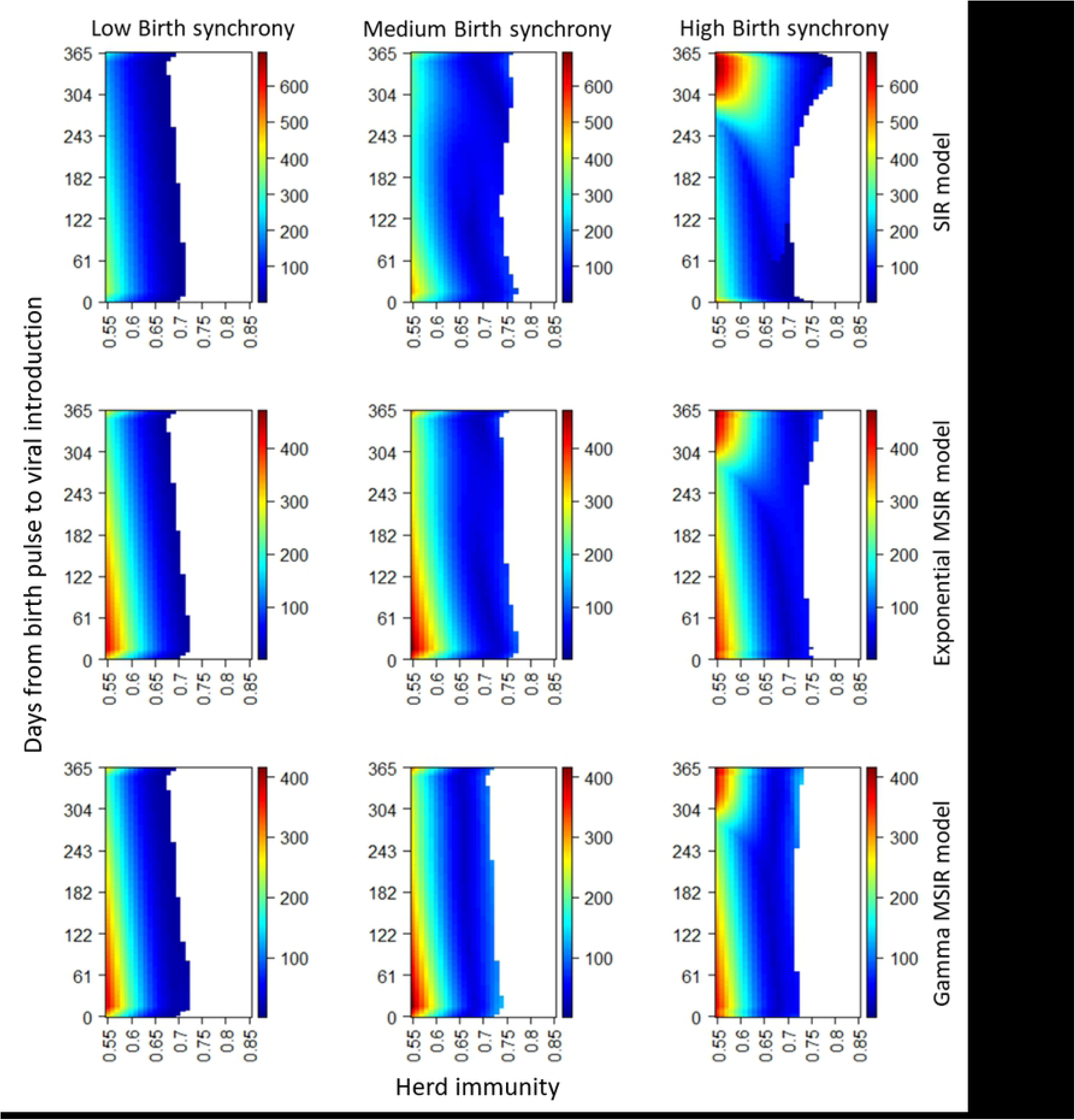
Magnitude of epidemics. The colour key shows the maximum number of infected individuals in the epidemic following viral introduction into a population. Herd immunity indicates the proportion of recovered individuals in a population, and days from birth pulse to viral introduction refers to the days of viral introduction since the peak of the previous birth pulse. Top, middle, and bottom panels show the results from SIR, exponential MSIR, and gamma MSIR models, respectively. Left, centre, and right panels show the results from low, medium, and high birth synchrony (s = 0.013, 1.3, and 130), respectively. White parts indicate no epidemic.

### Viral fadeout and persistence

We examined the effect of herd immunity and number of days from birth pulse to viral introduction on viral persistence. Introduction of infected individuals did not result in maintenance of infections in the population when birth synchrony was low (s=1.3) or when synchrony was high (s=130) but there was no MatAb (**Error! Reference source not found.**). Medium birth synchrony (s=13) appeared optimal for viral persistence. With high birth synchrony, if viral introduction occurred in presence of many susceptible individuals, very large epidemics were followed by deep troughs. Otherwise, if viral introduction occurred in presence of only few susceptible individuals, epidemics were not triggered, let alone viral persistence. The mechanisms of viral fadeout were described in Supplementary Text S2.

**Fig. 2.**
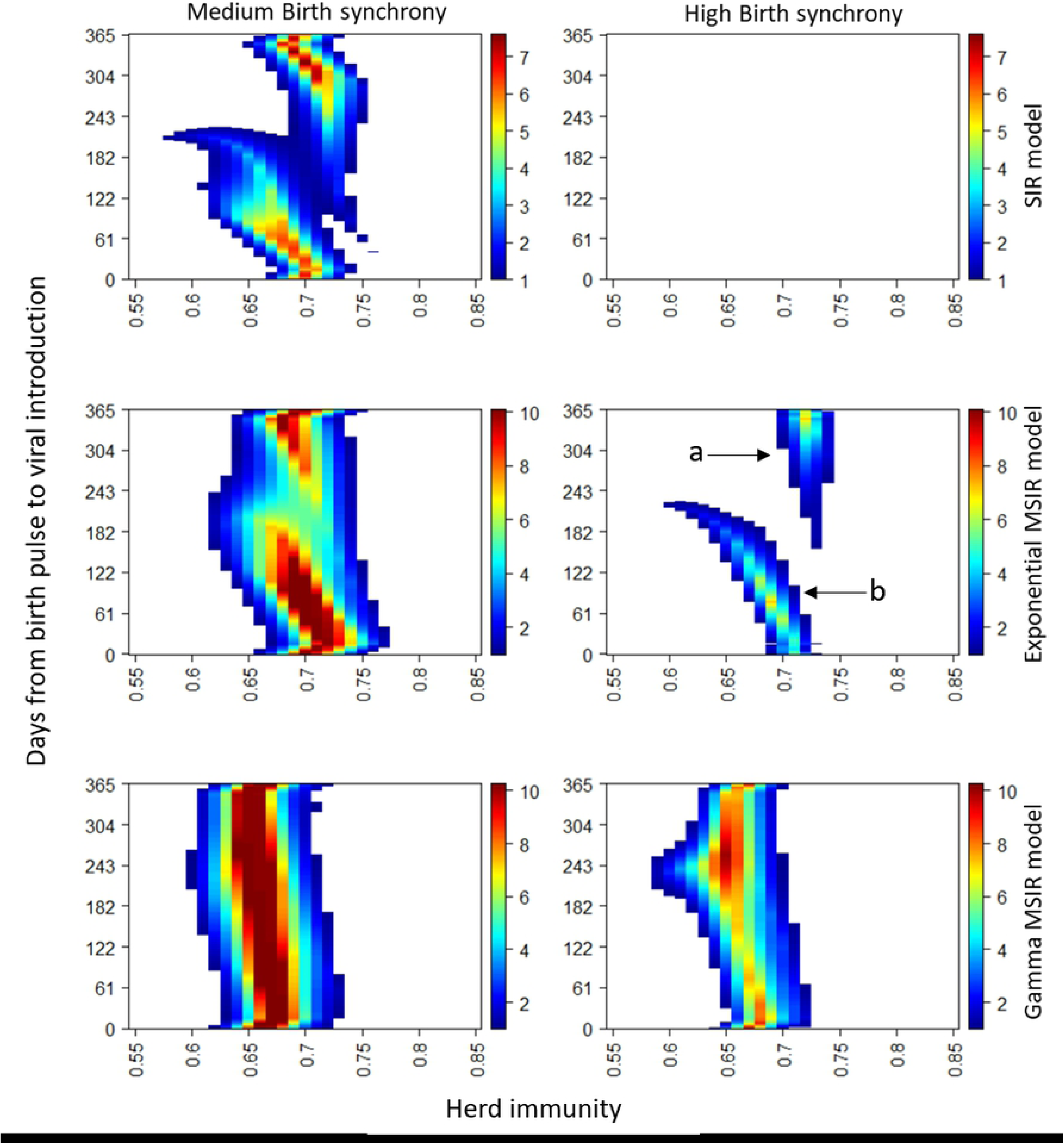
Number of infected individuals at the troughs following epidemics. Layout of panels is explained in Figure 1, but the results from low birth synchrony are not shown here. White parts indicate viral extinction.

Regarding viral persistence, the highest number of infected individuals at the troughs, following epidemic peaks, was found in two parts of the parameter space, for both small and large values of days from birth pulse to viral introduction (a and b in the middle panel of the right column in **Error! Reference source not found.**). The part ‘a’ was induced when a birth pulse correlated with decreasing prevalence, shortly after viral introduction; the part ‘b’ arose when a birth pulse correlated with decreasing prevalence after epidemic peaks. Compared to the exponential MSIR model, the gamma MSIR model showed more combinations of days from birth pulse to viral introduction and herd immunity that allowed viral persistence with high birth synchrony (s=130). This effect arose because gamma-distributed periods of maternally derived immunity resulted in the relatively steady supply of susceptible hosts throughout the year, just as medium birth synchrony (s=13) resulted in a steadier supply of susceptible hosts compared to high birth synchrony. Thus, the steady supply caused less intense epidemics and less deep (shallow) troughs.

### Timing of epidemic peaks

Overall, maternally-derived immunity delayed the timing of epidemic peaks. Maternally-derived immunity dispersed the supply of susceptible bats into the population, and thereby the number of infected bats relatively gradually increased, developing over a longer time period before peaking. When the timing of epidemic peaks was displayed in terms of months within a year, the SIR model showed seasonally clustered epidemic peaks, although this feature was weakened in the exponential MSIR model, and hardly observed in the gamma MSIR model (Fig. 3). When herd immunity was similar to herd immunity of an endemic equilibrium state in SIR model, the epidemic peaks mostly appeared within one year after viral introduction in the SIR model with birth pulses. However, in the exponential MSIR model, the epidemic peaks were delayed compared to those in the SIR model and showed a broader spectrum of timings than in the SIR model (Supplementary Fig. S3). In the gamma MSIR model, after viral introduction, the epidemics required a specific period to reach their peak, and the periods until epidemic peaks were delayed compared to those in the exponential MSIR model.

**Fig. 3.**
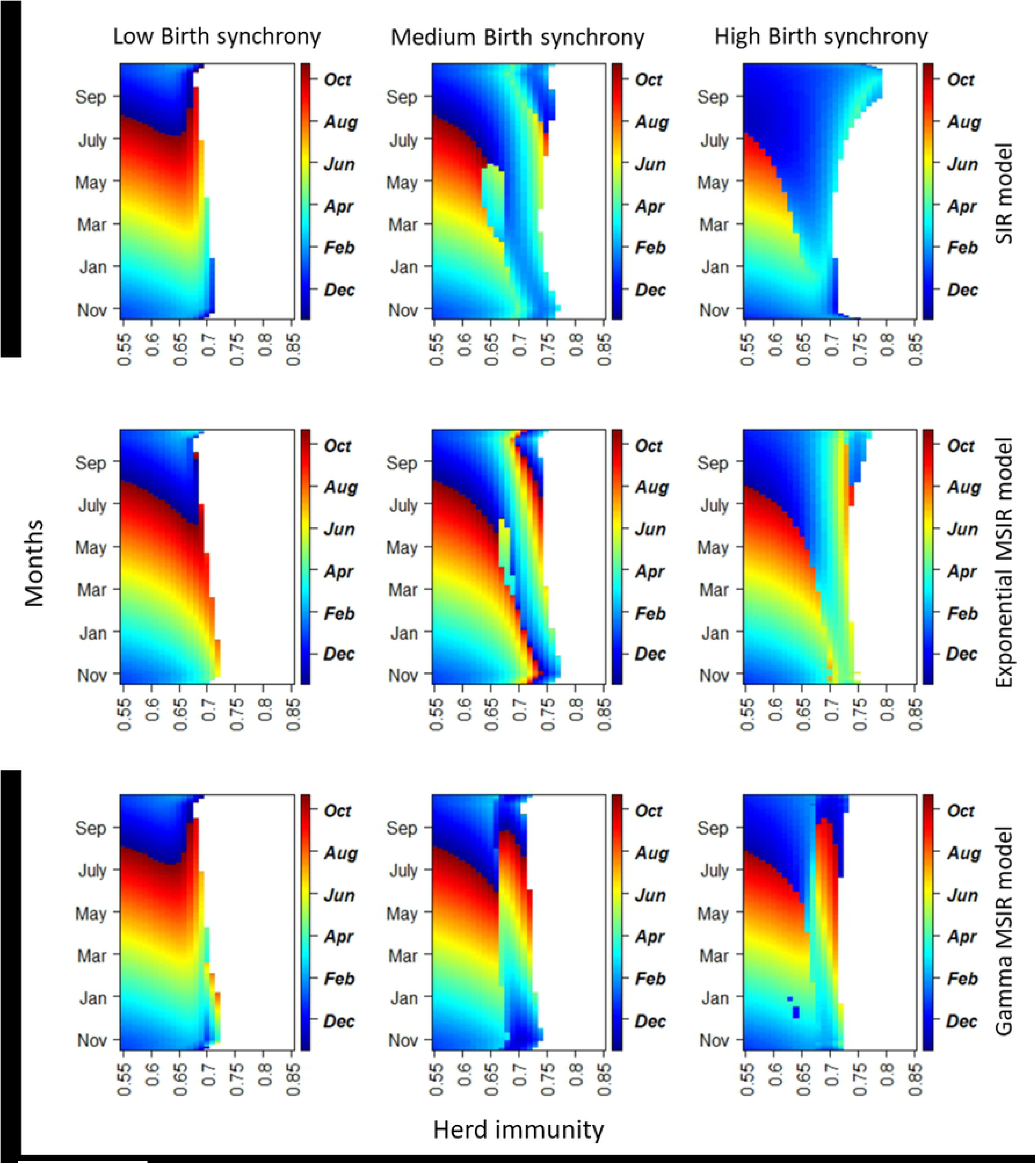
Timing of epidemic peaks. Peak of birth pulses occurred on 1st November. The colour key shows the month when the epidemics reach their peaks following viral introduction into a population. Layout of panels is explained in Fig. 1.

Taken together, in the exponential MSIR model, the epidemic patterns appeared to be more affected by seasonal birth pulses than by maternally derived immunity. In the exponential MSIR model, we modelled the number of maternally immune bats to decrease by as much as 1/255 of the remaining maternally immune bats every day. As a result, most juvenile bats lost their maternal immunity soon after birth, rather than 255 days after birth (Supplementary Fig. S2). When herd immunity was high and maternally immune newborns were more common than susceptible newborns, the temporal trend of epidemics was expected to change noticeably at 255 days after a birth pulse (the period of maternally derived immunity) rather than at the birth pulse (see Supplementary Fig. S4 for the number of individuals in each compartment across time). However, the number of infectious individuals began to rise at the time of birth pulses, which could probably be attributed to the exponential method of modelling the loss of maternally derived immunity. In comparison, the gamma MSIR model showed a more enhanced effect of maternally derived immunity (**Error! Reference source not found.**). In the gamma MSIR model, loss of maternally derived immunity seemed to occur mainly at 255 days after birth. Therefore, the impact of maternally derived immunity in determining epidemic pattern was much higher than in the exponential MSIR model. Moreover, the two relatively separate timings ensured a steady supply of susceptible individuals throughout the year, decreasing the extent of seasonal clustering of epidemic peaks. Another reason for birth pulses having a stronger impact on epidemics than did loss of maternally derived immunity was that the temporal synchrony of birth pulses was higher than that of loss of maternally derived immunity. Although more susceptible hosts were supplied from the loss of maternally derived immunity than from birth pulses, the tighter span of the latter relative to the former had a high impact on epidemic patterns.

**Figure 4.**
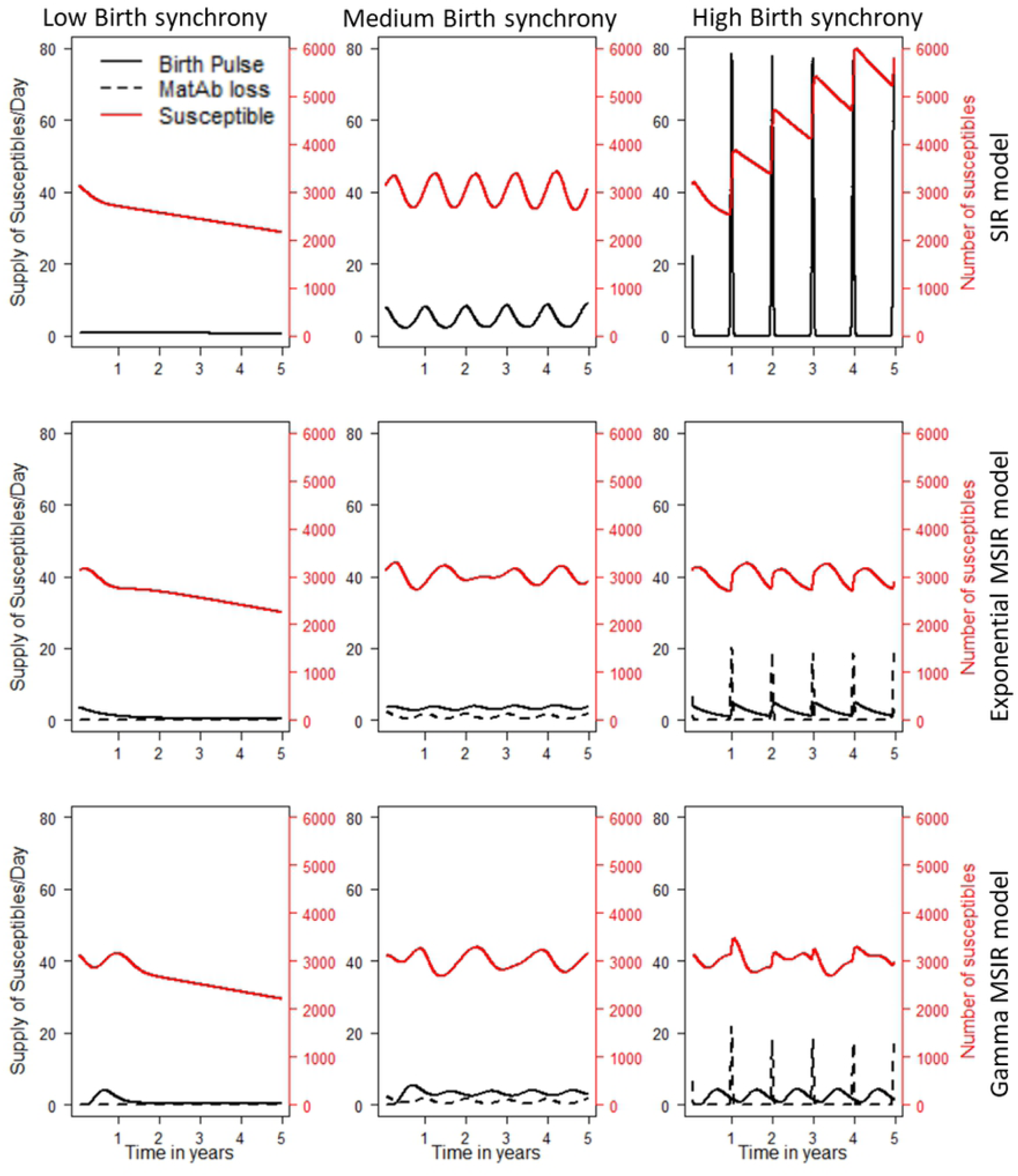
Supply of susceptible individuals from birth pulses and waning maternal immunity. Layout of panels is explained in Figure 1. The number of days since the last birth pulse before viral introduction until the viral introduction into a colony was 10, and herd immunity of the population was 0.7.

## Discussion

This study suggests that the effect of maternally-derived immunity in seasonal breeding species is not negligible and should be considered to improve our understanding of viral invasion and persistence in wildlife host populations. Modelling results showed that loss of maternally derived immunity disperses the timing of supply of susceptible individuals. The dispersed timing causes an increase in the likelihood of viral persistence and shifts the timing of epidemic peaks farther away from the timing of a birth pulse. However, these conclusions need to be interpreted with caution, since the modelling results are contingent on the assumptions made. In particular, SIR dynamics is one of at least three possible scenarios that describes Hendra virus dynamics [29]. Therefore, prediction of the timing of Hendra virus excretion pulses cannot be achieved by the results obtained from our models; instead, this study identified the relative importance of births and waning maternally derived antibody as the sources of susceptible hosts, and the significance of modelling methods of waning maternally derived antibody.

Seasonal clustering of Hendra virus spillover has been observed only in southeast Queensland and Northern NSW, whereas it occurred in northeast Queensland throughout the year [24]. While *P. alecto* in southeast Queensland and Northern NSW has a high birth synchrony, that in a more northern area has an inconsistent birth synchrony [32]. The modelling results showed medium birth synchrony to result in less temporal clustering of epidemic peaks than high birth synchrony (Fig. 3). The different temporal trend of spillover could, therefore, be associated with different synchrony of birth pulses of *P. alecto* based on latitude. Other climatic or ecological factors may also contribute to the different timing of spillover events (Peel et al. 2017) [23].

As flying foxes are natural reservoir hosts of Hendra virus [41], the virus must be maintained within the flying fox populations. However, seasonal birth pulses make it more difficult for the virus to persist compared to a population that has a constant birth rate [1]. Furthermore, when the annual birth pulse is tight, the disadvantage is further intensified [3]. For seasonally breeding flying foxes, other mechanisms are required to overcome or offset the disadvantageous conditions for viral persistence. Our modelling results support the implication that MatAb can play a role in mitigating adverse conditions for the maintenance of non-pathogenic virus, by dispersing the timing of supply of susceptible hosts into the population [42]. This is consistent with recent stochastic models showing that MatAb supports the maintenance of a henipavirus and lyssavirus in seasonally breeding African fruit bats (*Eidolon helvum*) [7, 42].

Wearing, Rohani (12) demonstrated that infectious disease modelling-based predictions are significantly affected by whether the infectious and latent periods are modelled using an exponential distribution or a gamma distribution. Heterogeneity in the longevity of protective immunity in different individuals requires mechanisms to deal with the distribution of immune periods of individuals in a model structure [43]. Here, we showed that gamma-distributed periods of maternally-derived immunity could generate significantly different modelling results compared to exponentially distributed periods. Since the functional form of the loss of MatAb against Hendra virus or against other viruses is unknown, we set g=8 to illustrate the effect of reducing the coefficient of variation in the duration of immunity by a third, namely from 100% (exponential distribution) to about 35% 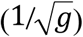), trying to realistically imitate waning MatAb. Results of the gamma MSIR model should not be interpreted as demonstrating a more accurate prediction than that by the exponential MSIR model; it merely shows that substantially different modelling results can be generated depending on whether the loss of maternally-derived immunity is exponentially or gamma-distributed. Therefore, appropriate modelling of when maternally immune newborns lose their passive immunity, depending on species and pathogens against emerging infectious diseases is expected to help improve the prediction of disease outbreaks.

Although this study modelled epidemics in a single population, the results obtained should be considered in the context of metapopulation structure, since this study assumed SIR dynamics in which metapopulations would be expected to play an important role in Hendra virus maintenance [29, 30]. An effect of the seasonality of epidemics in metapopulations is that seasonal forcing of transmission may cause the epidemics in each population to be synchronised [44]. Synchronised epidemics are likely to increase the probability of viral fadeout in a metapopulation. Viral introduction would be less probable if epidemics in populations become extinct at similar times. It is, therefore, necessary to examine the effects of seasonality of viruses on the synchrony of epidemics in the metapopulation models of hosts.

Given the growing interest in emerging zoonotic diseases from wildlife [45], there is a need to combine empirical studies of zoonotic reservoir ecology with mathematical models to develop a mechanistic understanding of virus persistence and spillover. By incorporating maternally derived immunity into generic epidemic models, we have provided a framework to study epidemics in seasonally breeding wildlife species. Complex effects of demographic and virus-related parameters have previously been reported in other systems [5]. We explored a range of plausible assumptions to complement the limited empirical data available for the dynamics of Hendra virus in flying fox populations, and the results showed the effects of maternally derived immunity on the timing of epidemic peaks, rather than predicting the actual timing of epidemic peaks. Caution must be exercised, however, when arriving at conclusions concerning the effect of maternally derived immunity, given the specific circumstances associated with each disease and host population structure.

## SUPPORTING INFORMATION

Additional Supporting Information may be found in the online version of this article:

## Acknowledgements

We thank J. McBroom, L. Grogan, D. Kerlin, K. Wells, and J. Giles for assistance in analysing data and structuring models. We thank Stephanie Palmer for editing this manuscript.

